# Fatty acid starvation activates RelA by depleting lysine precursor pyruvate

**DOI:** 10.1101/635748

**Authors:** Anurag Kumar Sinha, Kristoffer Skovbo Winther, Mohammad Roghanian, Kenn Gerdes

## Abstract

Bacteria experiencing nutrient starvation induce the ubiquitous stringent response, a profound physiological change that reprograms cell physiology from fast to slow growth and stress survival. The stringent response is mediated by the secondary messengers pppGpp and ppGpp collectively referred to as (p)ppGpp or “alarmone”. In *Escherichia coli*, two paralogs, RelA and SpoT, synthesize (p)ppGpp. RelA is activated by amino acid starvation whereas SpoT, which can also degrade (p)ppGpp, responds to fatty acid (FA), carbon and phosphate starvation. Here, we discover that FA starvation leads to rapid activation of RelA and reveal the underlying mechanism. We show that fatty acid starvation leads to depletion of lysine that, in turn, leads to the accumulation of uncharged tRNA^lys^ and activation of RelA. SpoT was also activated by fatty acid starvation but to a lower level and with a delayed kinetics. Next, we discovered that pyruvate, a precursor of lysine, is depleted by FA starvation. We also propose a mechanism that explains how FA starvation leads to pyruvate depletion. Together our results indicate that many responses to nutrient depletion may ultimately result indirectly from depletion of amino acids and thereby activating RelA. Interestingly, FA starvation provoked a ~100-fold increase in *relA* dependent ampicillin tolerance.

## Introduction

Bacteria growing in the environment often experience limiting accessibility of nutrients. Under such conditions, bacteria mount the stringent response that alters the levels of a wide range of gene transcripts and metabolites (Traxler *et al.*, 2008, Durfee *et al.*, 2008, Sanchez-Vazquez *et al.*, 2019). These global changes are controlled by the secondary messengers pppGpp and ppGpp, the effector molecules of the stringent response collective called (p)ppGpp or “alarmone”. Importantly, (p)ppGpp is required for almost all pathogenic bacteria to be virulent and in a number of cases also for expressing antibiotic tolerance (Dalebroux *et al.*, 2010, Harms *et al.*, 2016). In addition, the levels of (p)ppGpp vary inversely with bacterial growth-rates during balanced growth, allowing for adjustment of cellular growth-parameters also in the absence of external stress (Potrykus *et al.*, 2011).

In *E. coli*, two paralogous enzymes, RelA and SpoT, are responsible for the metabolism of (p)ppGpp (Ronneau & Hallez, 2019). RelA has (p)ppGpp synthetase activity whereas SpoT has both synthetase and hydrolase activities. During amino acid starvation, RelA associates with uncharged tRNA and this binary complex is loaded at “hungry” ribosomal A-sites thereby leading to activation of RelA (Brown *et al.*, 2016, Loveland *et al.*, 2016, Arenz *et al.*, 2016, Winther *et al.*, 2018, Kushwaha *et al.*, 2019). Accumulation of (p)ppGpp leads to major changes in bacterial physiology: transcription of rRNA and tRNA operons is strongly reduced, whereas transcription of amino acid biosynthetic operons is stimulated to replenish amino acids pools (Durfee *et al.*, 2008, Traxler *et al.*, 2008). High (p)ppGpp levels thus lead to slow growth or growth arrest. After readjusting cellular physiology, SpoT hydrolyses (p)ppGpp and thereby allowing resumption of cell growth at an adjusted rate (Potrykus & Cashel, 2008)*. *ΔrelA* ΔspoT* double mutant strains (also called (p)ppGpp^0^ strains) are unable to synthesize (p)ppGpp and die in minimal media but can grow in rich media (Xiao *et al.*, 1991). Such strains exhibit auxotrophy for multiple amino acids, survive less well during stationary phase and exhibit aberrant cell morphology (Hauryliuk *et al.*, 2015, Dalebroux *et al.*, 2010).

While the mechanism of RelA activation during aa starvation is well understood (Kushwaha *et al.*, 2019, Winther *et al.*, 2018), it is more unclear as to how other starvation conditions lead to the accumulation of (p)ppGpp. A number of studies implicate SpoT in (p)ppGpp accumulation during multiple stress conditions, including fatty acid (FA) starvation, either by reducing SpoT-dependent (p)ppGpp-hydrolase or by increasing (p)ppGpp-synthetase activity, or both (Xiao *et al.*, 1991, Seyfzadeh *et al.*, 1993, Vinella *et al.*, 2005, Spira *et al.*, 1995, Wang *et al.*, 2016, Kim *et al.*, 2018). Inhibition of FA chain elongation by the drug cerulenin, which inhibits FabB/FabF (Fig. 1A), stimulates SpoT synthetase activity (Battesti & Bouveret, 2006, Seyfzadeh *et al.*, 1993). SpoT is believed to sense FA limitation by interaction with Acyl Carrier Protein (ACP), the central cofactor of FA synthesis (Battesti & Bouveret, 2006, Battesti & Bouveret, 2009). Remarkably, while Nomura and coworkers showed that SpoT synthesizes ppGpp under FA starvation. The contribution of RelA to synthesize ppGpp under this condition was not analysed (Seyfzadeh *et al.*, 1993).

**Figure 1.**
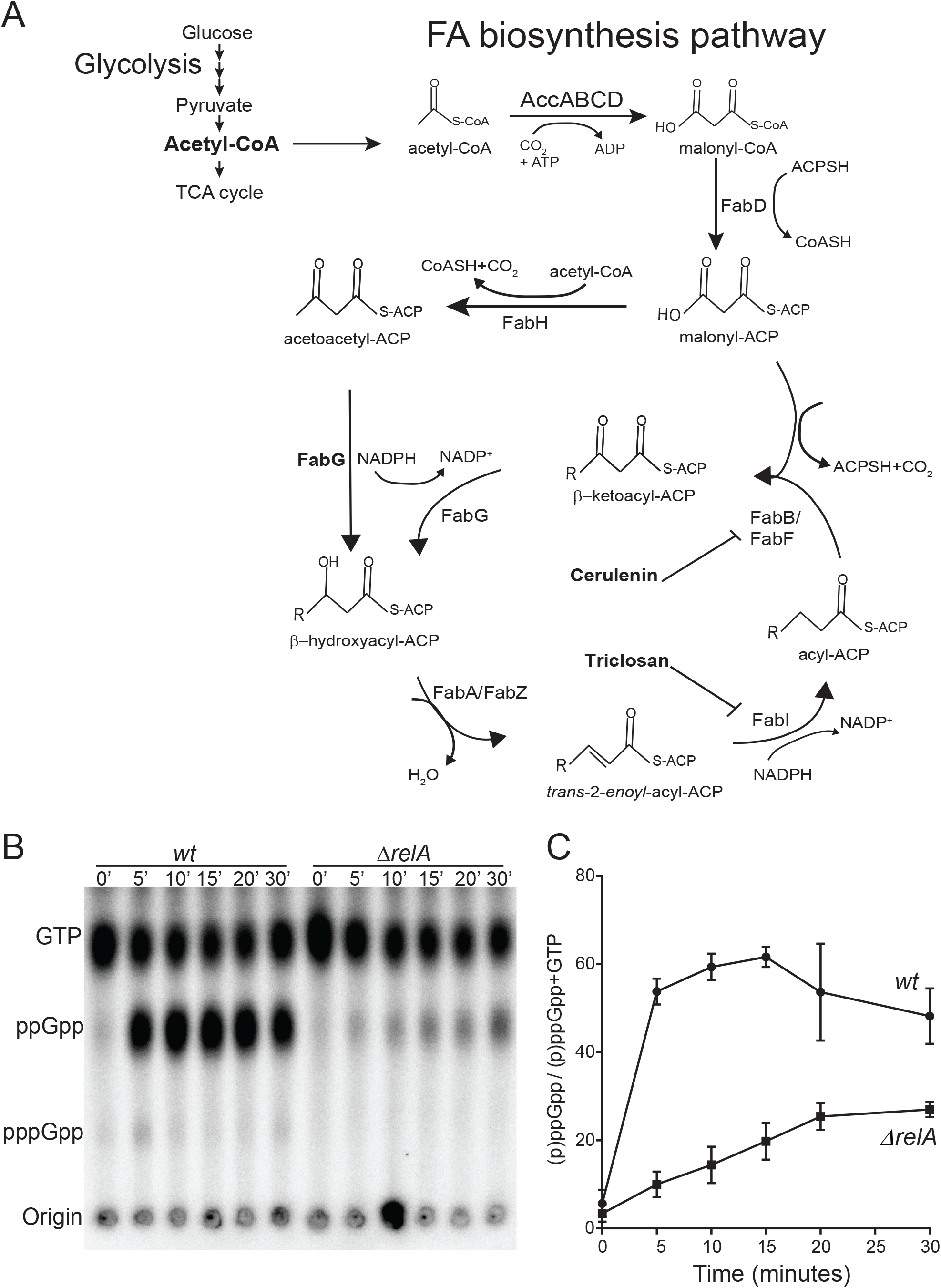
RelA is the major (p)ppGpp synthetase during fatty acid starvation. **(A)** Schematic representation of FA biosynthesis in *E. coli*. Acetyl-CoA is the basic unit for FA synthesis and is synthesized either from pyruvate by pyruvate dehydrogenase or generated by β–oxidation of FA. The first committed step in FA synthesis is catalyzed by the multi-subunit enzyme complex acetyl-CoA carboxylase (ACC) by carboxylation of acetyl-CoA to malonyl-CoA. The malonyl group is then transferred to acyl carrier protein (ACP) by FabD to form malonyl-ACP. The initial condensation reaction is performed by FabH that catalyzes the condensation of malonyl-ACP and acetyl-CoA to form the 4-C compound acetoacetyl-ACP. The acetoacetyl-ACP is substrate of cyclic reactions performed by FabG, FabA/FabZ and FabI to make 4-C acyl-ACP. FabB/FabF catalyse chain elongation by condensation of 4-C acyl-ACP with a new molecule of malonyl-ACP thereby increasing the chain length by 2-C in each cycle. As indicated, cerulenin and triclosan inhibit FA chain elongation by inhibiting FabB/FabF and FabI, respectively. **(B)** Kinetics of (p)ppGpp accumulation in MG1655 (*wt*) and Δ*relA* strains after the addition of cerulenin (200 µg/ml) revealed by TLC and radiolabeling with ^32^P as described in *Experimental Procedures* according to the protocol kindly provided by Mike Cashel (Cashel, 1994) with a few modifications (Winther *et al.*, 2018). Positions of GTP, ppGpp and pppGpp are indicated. Cells were grown in MOPS media supplemented with 0.2 % glucose at 37°. **(C)** Quantification of (p)ppGpp accumulation in *wt* and Δ*relA* cells. Quantification was done as described in *Experimental Procedures*. Data collected from 4-5 separate experiments were used to calculate mean with standard deviations. MIC for both *wt* and *ΔrelA* cells were found to be close to ~64 µg/ml for cerulenin and no growth observed after 24 hrs of incubation (Fig. S1A). Initially, we used 200 µg/ml cerulenin (~4X MIC) to inhibit FA synthesis for (p)ppGpp measurements. This concentration was also used in previous studies (Battesti & Bouveret, 2006, Seyfzadeh *et al.*, 1993).

Here we show that inhibition of FA synthesis in *wt* cells causes rapid accumulation of uncharged lysine tRNA, activation of RelA and synthesis of (p)ppGpp. Consistently, addition of lysine to the growth media abolishes RelA activation after FA starvation. Addition of pyruvate, an essential precursor of lysine and FA synthesis significantly reduces RelA activation. Interestingly, under FA starvation, the *wt* strain exhibited an almost 100-fold increased level in antibiotic tolerance as compared to the *ΔrelA* mutant cells, an effect that was abolished by the addition of lysine. Based on these results, we propose that increased consumption of pyruvate under FA starvation leads to lysine starvation and activation of RelA that, in turn, explains the increased antibiotic tolerance.

## Results

### Fatty acid starvation induces rapid RelA-dependent accumulation of ppGpp

We measured (p)ppGpp accumulation after treating cells of strain MG1655 (*wt*) and its *ΔrelA* derivative with cerulenin. In the *wt* strain, we saw an immediate and strong increase in the level of ppGpp whereas the response of the *ΔrelA* strain was much slower and at no point reached the level of the *wt* strain (Fig. 1B). Only ppGpp accumulated since almost no pppGpp was observed after cerulenin treatment. Quantification of the response revealed that ppGpp level reached ≈60% of total labelled nucleotides (ppGpp + GTP) within 5 minutes in the *wt* strain, and then decreased to a level of 40-50% (Fig. 1C). In the Δ*relA* strain, the [ppGpp] increased very slowly with less than 10% ppGpp accumulated in the first 5 minutes, increasing slowly up to ~25 % in 30 minutes (Fig. 1C). These results show that RelA is responsible for the rapid ppGpp accumulation during FA starvation. It should be noted that *wt* and *relA* strains exhibited similar MICs toward cerulenin, excluding the possibility that the reduced level of ppGpp in the *relA* strain was due to different sensitivity of the cells (Fig. S1A).

### Inhibition of fatty acid synthesis induces lysine starvation

Addition of all 20 amino acids to *wt E. coli* cells growing in minimal medium prevented ppGpp accumulation during cerulenin treatment, indicating that inhibition of FA synthesis induces amino acid starvation (Seyfzadeh *et al.*, 1993). To further analyse this phenomenon, we measured ppGpp accumulation in the presence of polar (Mix A) or non-polar amino acids (Mix B) (Fig. 2A).

**Figure 2.**
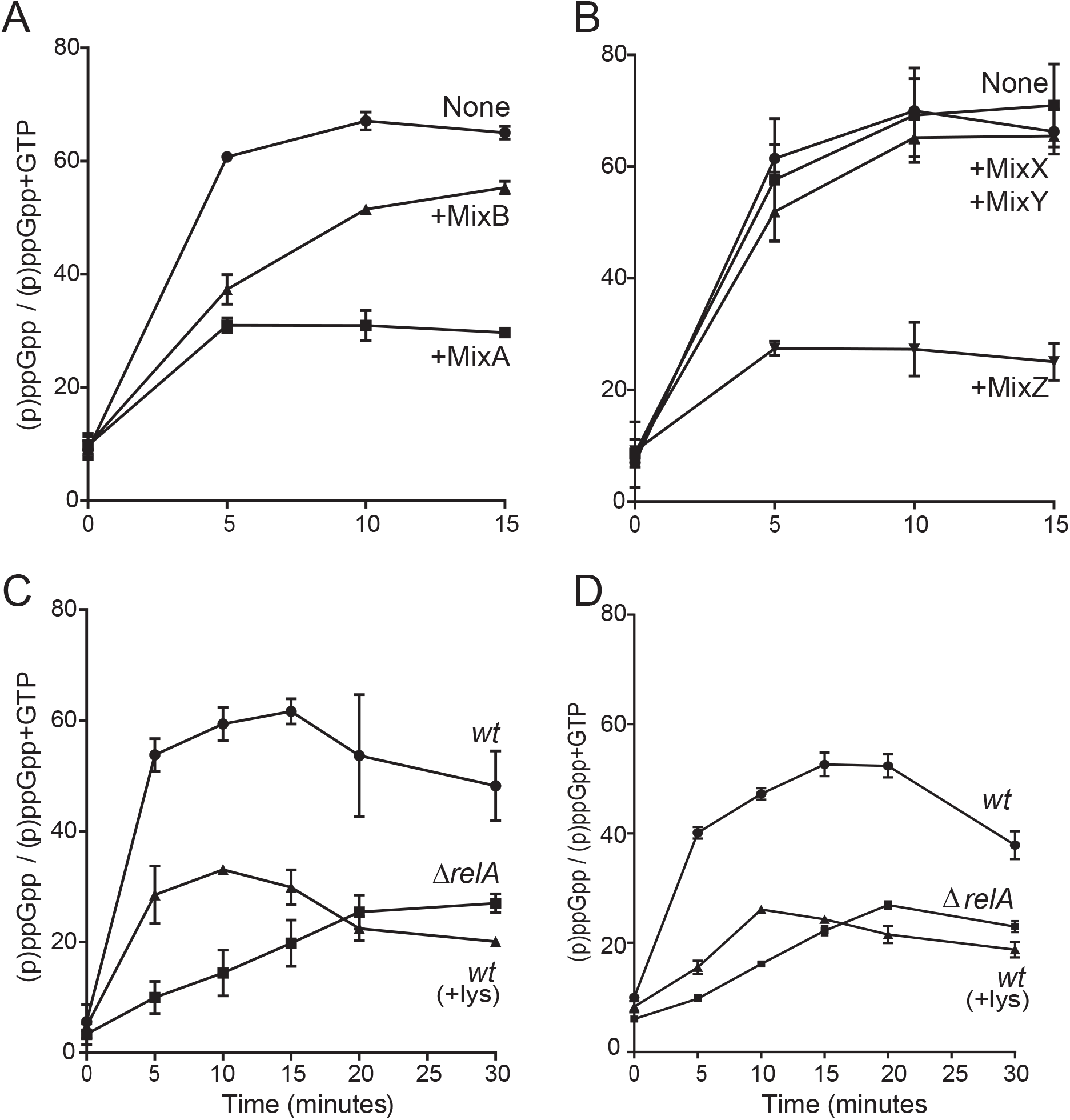
Exogenous lysine suppresses activation of RelA during fatty acid starvation. **(A)** Deconvolution of aa depletion by supplementing the MOPS glucose media at 37° with aa mixtures 10 minutes before the addition of cerulenin (200 µg/ml). MixA: alanine, arginine, asparagine, aspartate, glutamate, glutamine, glycine, histidine, serine and lysine); MixB (isoleucine, proline, tyrosine, threonine, tryptophan, valine, leucine, cysteine, methionine and phenylalanine). Final aa concentrations were as recommended (Neidhardt *et al.*, 1974). **(B)** MixA was subdivided into 3 sub-mixes: MixX (alanine, arginine, asparagine), MixY (aspartate, glutamate, glutamine) and MixZ (glycine, histidine, serine and lysine). These mixtures were supplemented separately into different *wt* cultures and then the (p)ppGpp levels were measured before and after the addition of cerulenin (200 µg/ml). **(C)** and **(D)** Quantification of (p)ppGpp accumulation in *wt*, Δ*relA* and *wt* (+lys; supplemention with 0.4 mM lysine) upon high (200 µg/ml) or low (50 µg/ml) cerulenin concentration, respectively. The quantifications shown were derived from at least three different experiments as mean values with standard deviations, and primary data were obtained as described in the Legend to Fig. 1B.

Addition of Mix A drastically reduced ppGpp accumulation upon cerulenin treatment (Figs 2A and S2A). Mix A was sub-divided into three groups: Mix X (alanine, arginine and asparagine), Mix Y (aspartate, glutamate and glutamine) and Mix Z (glycine, histidine, serine and lysine). Supplementation of mix Z decreased ppGpp accumulation similar to mix A, whereas addition of mix X and mix Y had no effect (Figs 2B and S2B). Further deconvolution of Mix Z revealed that lysine alone was sufficient to suppress RelA activation (Figs 2C and S2C-D). We observed that 0.4mM lysine supplementation unable to completely suppress RelA activation at early time points (5 and 10 minutes time points) probably due to high uncharging of lysine tRNA (shown below). We could not supplement higher amount of lysine as at higher amount it feedback inhibits lysC, first common enzyme for synthesis of methionine, threonine and lysine. The above experiments were accomplished using a high concentration of cerulenin (200 µg/ml; ~4X MIC). However, lysine supplementation abolished RelA dependent ppGpp accumulation also at a low concentration of cerulenin (50 µg/ml; ~1X MIC; Fig. 2D).

To investigate FA inhibition in general leads to lysine starvation, we analysed the response to triclosan that inhibits FA synthesis by targeting FabI (Fig. 1A) (Heath & Rock, 1999). Similar to cerulenin treatment, *wt* cells accumulated more than 60 % of ppGpp within 5 minutes of triclosan addition and then slowly decreased to ~40 % in ~30’ (Fig. S3). In the Δ*relA* strain, the ppGpp level increased much slower and was in the range of 10-15 % of that of the wt, very similar to the result with cerulenin (Fig. 1B and 1C). Again, supplementation of lysine curtailed ppGpp accumulation in the *wt* strain, and after 15 minutes of treatment, the level was comparable to that of the Δ*relA* strain (Figs S3A and S3B). In summary, our results show that inhibition of FA synthesis depletes the lysine pool that, in turn, activates RelA to synthesize ppGpp.

### Fatty acid starvation induces accumulation of uncharged tRNA^Lys^

To further substantiate this conclusion, we analysed the charging levels of tRNA^lys^ by Northern blotting analysis. In *wt* cells, the level of uncharged tRNA^lys^ increased from 12% before treatment to 72%, 5 minutes after triclosan treatment (Fig. 3A). Addition of lysine reduced uncharged tRNA^lys^ levels to only 09%, similar to before starvation (Fig. 3A). Next, we analysed the effect of FA starvation on the charging level in a Δ*relA* strain. Here, a relatively high level of uncharged tRNA^lys^ was present already before starvation (22%) that increased to 78% after 5 minutes with triclosan. Similar to *wt* cells, lysine supplementation reduced uncharged tRNA^lys^ to 27%. (Fig. 3A).

**Figure 3.**
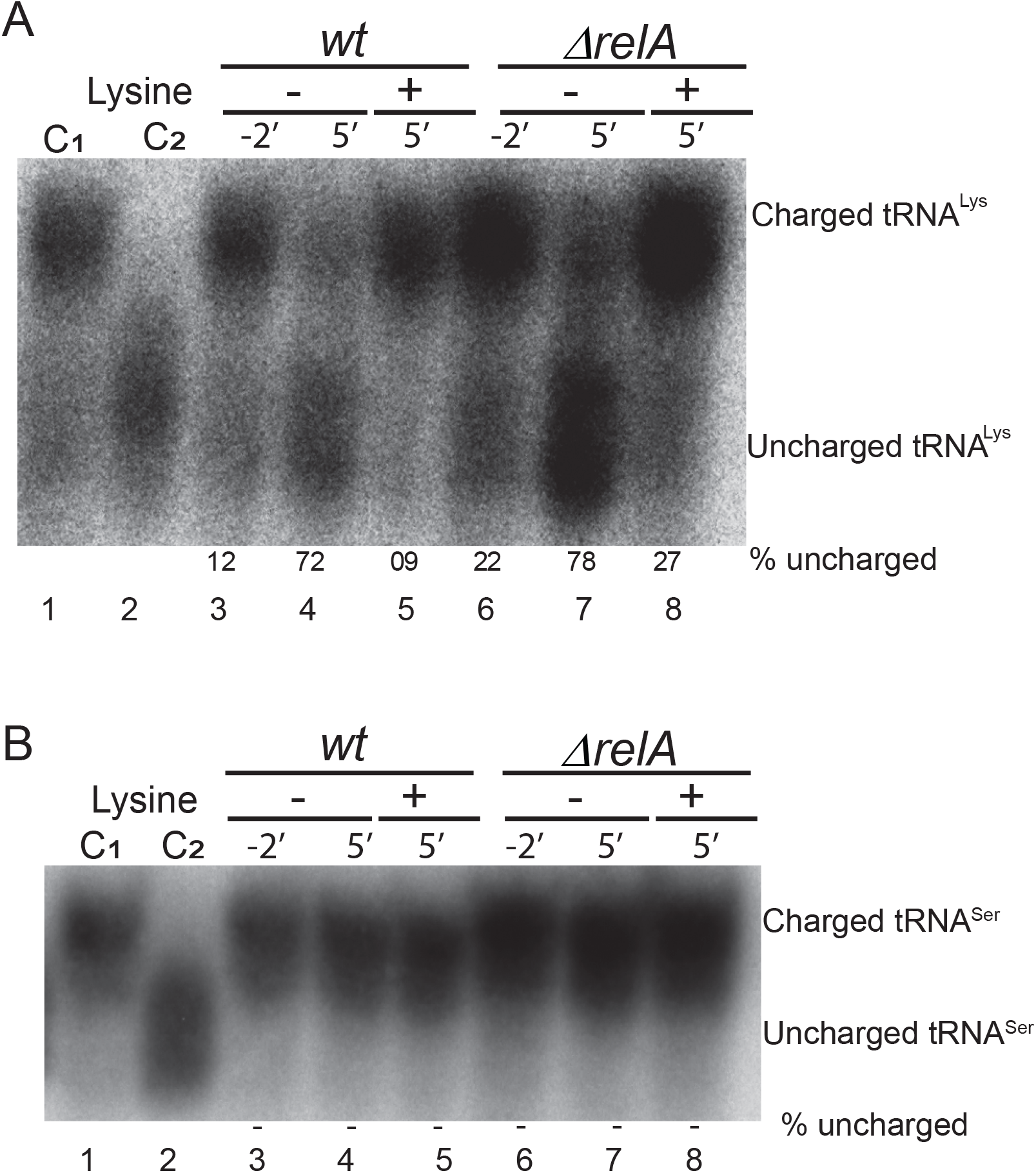
Lysine curtails accumulation of uncharged tRNA^lys^ after fatty acid starvation. (**A**) Charging levels of tRNA^lys^ before and after the addition of triclosan (1µg/ml). Total RNA was separated on PAGE, processed for Northern blotting analysis and hybridized with a probe specific for tRNA^lys^ (*Experimental Procedures*). Representative image shown here. Percentage (%) of uncharged tRNA was quantified according to the signal intensity of uncharged/(charged+uncharged). One culture was supplemented with lysine before triclosan addition. A fraction of tRNA was deacylated *in vitro* by base treatment (100 mM Tris-HCl [pH-9]) for control (C_2_) and indicates the position of uncharged tRNA^lys^. C_1_-Charged tRNA control. (**B**) The filter from (A) was stripped and hybridized with a tRNA^ser^ specific probe.

We also monitored the kinetics of uncharging of tRNA^lys^ after triclosan treatment both in *wt* and in Δ*relA* cells (Fig. S4A & C). Interestingly, in Δ*relA* cells, level of uncharged tRNA^lys^ increased up to ~70%, 5 minutes after triclosan treatment and remained uncharged until 15 minutes (Fig. S4C). However, in *wt* cells, level of uncharged tRNA^lys^ increased up to ~70% at 5 minutes after triclosan treatment and then started decreasing at 10 minutes time point and went back to before starvation level at 15 minutes after treatment (Fig. S4A). This indicates that RelA mediated ppGpp produced in *wt* cells acts quickly to replenish the lysine pool by adjusting cellular physiology, which does not occur efficiently in Δ*relA* strain.

To investigate of the effect of FA starvation was specific to tRNA^lys^, we also analysed tRNA^ser^. Indeed, we found no accumulation of uncharged tRNA^ser^ (Fig. 3B and Fig. S4B & D). In conclusion, FA starvation specifically depletes the lysine pool, resulting in specific and high increase in the level of uncharged tRNA^lys^.

### Pyruvate prevents depletion of lysine during fatty acid starvation

To investigate the mechanism of lysine depletion, we thought possible that FA starvation could either provoke lysine degradation or inhibit lysine biosynthesis. The major pathway of lysine degradation occurs through its decarboxylation by two lysine decarboxylases: CadA and LdcC (Keseler *et al.*, 2017, Reitzer, 2005). To test first possibility, we analysed a *cadA ldcC* strain during FA starvation. However, the double deletion strain accumulated ppGpp similar to *wt* when treated with cerulenin, showing that degradation is probably not responsible for observed lysine depletion (Fig. S5).

The alternative possibility is that FA starvation inhibits lysine biosynthesis. In *E. coli*, lysine is synthesized from aspartate (Keseler *et al.*, 2017). Interestingly, two initial steps for the lysine, threonine and methionine biosynthesis pathways are common and they all utilize common substrate called aspartate 4-semialdehyde. The first unique step, also considered as the rate-limiting step, for lysine biosynthesis, is catalyzed by DapA (4-hydroxy-tetrahydrodipicolinate synthase). DapA catalyzes condensation of aspartate-4-semialdehyde and pyruvate to generate (2S,4S)-4-hydroxy-2,3,4,5-tetrahydrodipicolinate (Laber *et al.*, 1992) (Fig. 4A). Since, FA starvation only affects lysine and not aspartate, threonine or methionine, we exclude the possibility of any major change in the level of aspartate 4-semialdehyde. The other substrate of DapA is pyruvate. Pyruvate is also needed for synthesis of acetyl-CoA, the precursor of FA biosynthesis (Fig. 1A). Interestingly, previous reports showed FA starvation causes reduction in acetyl-CoA levels (Heath & Rock, 1995, Furukawa *et al.*, 1993). Therefore, we hypothesized that, during FA starvation, pyruvate will be utilized to replenish acetyl-CoA and thus will not be available for DapA to synthesize lysine.

**Figure 4.**
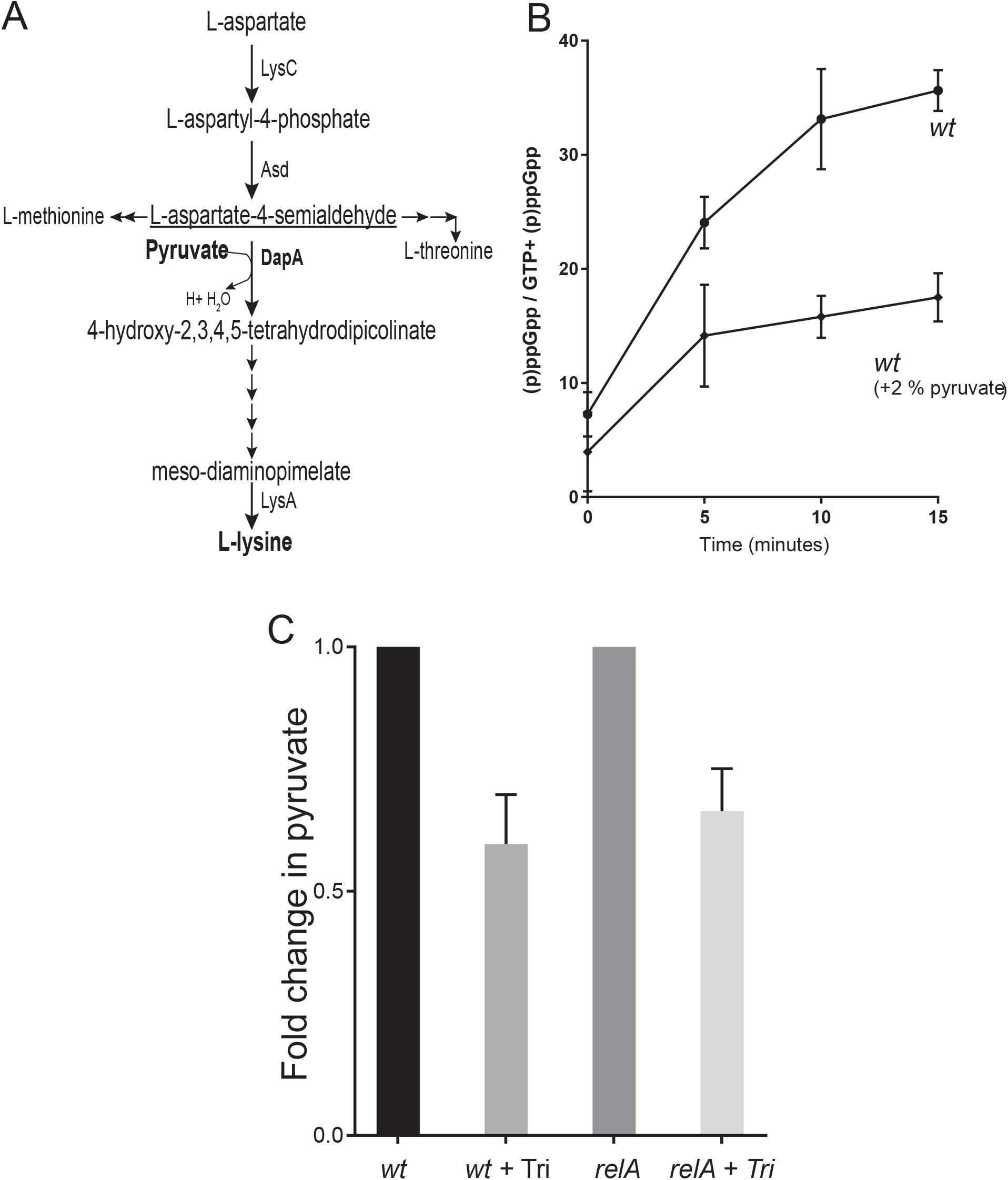
Pyruvate reduces accumulation of (p)ppGpp. **(A)** Schematic of the lysine biosynthesis pathway with relevant intermediates. Lysine is synthesized from aspartate in nine reactions. The initial two steps are common to lysine, methionine and threonine biosynthesis. The first unique and rate limiting step is catalyzed by DapA, which uses aspartate-4 semialdehyde and pyruvate to synthesize 4-hydroxy-tetrahydrodipicolinate. **(B)** Quantification of (p)ppGpp measurement of MG1655 (*wt*) cells, grown in MOPS + glycerol (0.5 %) medium at 37°C, supplemented with the 2% of pyruvate added before cerulenin (200 µg/ml) treatment. **(C)** Quantification of cellular levels of pyruvate before and after triclosan (Tri; 1µg/ml) treatment. Data presented here are the mean values from three different experiments.

To test this hypothesis, we grew cells of the *wt* strain in MOPS minimal medium supplemented with pyruvate. Indeed, addition of 2% pyruvate dramatically reduced the (p)ppGpp level (Fig. 4B and Fig. S6A). In this experiment, glycerol was used as carbon source (instead of glucose) because transport of pyruvate is more efficient in glycerol media (Kreth *et al.*, 2013). This led to a somewhat lower rate of (p)ppGpp accumulation (compare Figs 1C and 4B). Pyruvate-dependent suppression of RelA activation is specific to FA starvation as pyruvate addition only slightly affected RelA activation during induced isoleucine starvation (Fig. S6B).

We then directly quantified cellular pyruvate levels upon triclosan treatment. Remarkably, triclosan treatment immediately reduced cellular pyruvate levels up to 40-50 % compared to pre-treatment levels in both *wt* and in Δ*relA* strains (Fig. 4C), further substantiating our hypothesis that FA starvation leads to depletion of pyruvate.

### Fatty acid starvation induces ppGpp-dependent ampicillin tolerance

High cellular levels of (p)ppGpp induce antibiotic tolerance in *E. coli* (Dalebroux *et al.*, 2010, Rodionov & Ishiguro, 1995, Kusser & Ishiguro, 1985, Rodionov & Ishiguro, 1996). Therefore, we analysed antibiotic tolerance after treatment with cerulenin/triclosan. The *wt* strain exhibited an almost 100-fold increased ampicillin tolerance as compared to the *ΔrelA* strain when starved for FA (Fig. 5A and 5B). Ampicillin tolerance was abolished when lysine was added to the growth medium (Fig. 5A and 5B). Addition of valine that, in *E. coli* K-12, induces isoleucine starvation, also induced *relA*-dependent ampicillin tolerance (Fig. 5C), confirming earlier observations (Kusser & Ishiguro, 1985). Addition of higher doses of cerulenin (200 µg/ml; ~4X MIC) made almost all cells refractory to ampicillin killing both in *wt* and in Δ*relA* cells (Fig. S7A and S7B). These results show that activation of RelA by low levels of FA starvation is sufficient to confer antibiotic tolerance and that such tolerance can be eliminated by lysine supplementation. Thus, the ppGpp-mediated ampicillin tolerance cannot simply be explained by slow growth, as cerulenin treatment induced similar growth arrest in both *wt* and Δ*relA* strains at low concentration of cerulenin (50 µg/ml; ~1X MIC; Fig. S7C).

**Figure 5.**
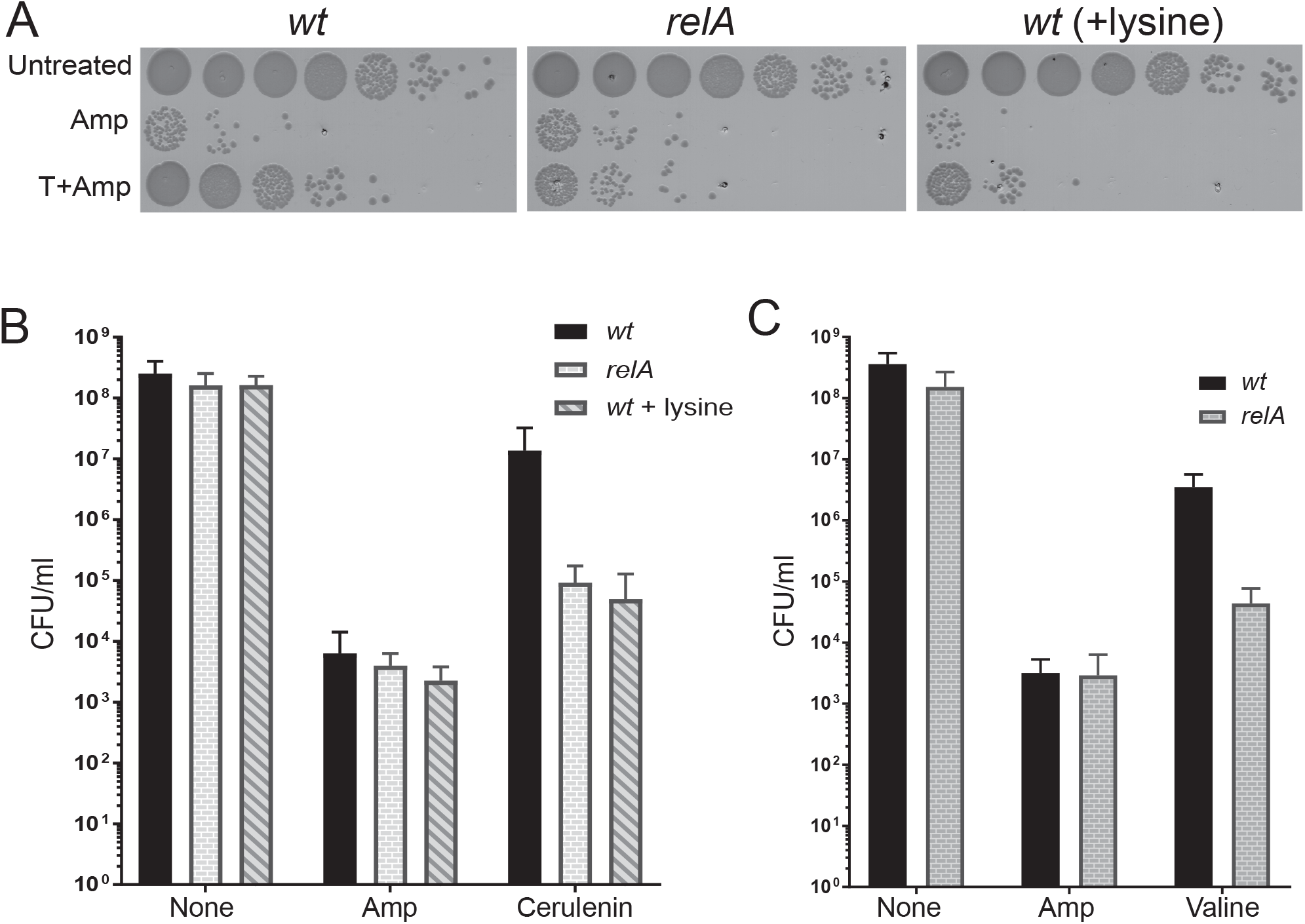
Fatty acid starvation induces ampicillin tolerance. **(A)** Cells of MG1655 (*wt*) and Δ*relA* strains growing balanced in MOPS + 0.2 % glucose media at 37°C were either left untreated, directly exposed to ampicillin (100 µg/ml) for 3 hrs or first treated with triclosan (1µg/ml ~MIC) for 5 minutes and then exposed to ampicillin. In another culture of *wt* cells, 0.4 mM lysine was added. **(B)** Averages of colony forming units (CFU) from four different experiments, performed in a same way as in (A) but treated instead with cerulenin (50 µg/ml). Mean and standard deviations are shown. **(C)** Effect of valine. Experiments were accomplished as in (B) except that the stringent response was induced by the addition of valine (500 µg/ml). Quantification from three experiments are shown here. Abbreviations: T, Triclosan (1µg/ml); C(50), Cerulenin (50 µg/ml); Val(500), Valine (500 µg/ml).

## Discussion

Here we show that RelA is the major contributor to the accumulation of (p)ppGpp during FA starvation. As described previously (Seyfzadeh *et al.*, 1993), SpoT also contributes, but by a significantly lower level (Fig. 1). This unexpected observation led us to test if supplement of exogenous amino acids would prevent activation of RelA. Indeed, addition of lysine abrogated RelA activation (Fig. 2). Consistently, FA starvation of the *wt* strain increased the ratio of uncharged to charged tRNA^lys^ (Fig. 3 and S4A). FA starvation of the *ΔrelA* strain led to an even stronger and persistent increase of the level of uncharged tRNA (Fig. 3A, right panel), consistent with the notion that (p)ppGpp curtails translation and consumption of aa during starvation (Svitil *et al.*, 1993, Rojas *et al.*, 1984).

FA starvation leads to a rapid and strong reduction of the level of pyruvate (Fig. 4C). Since pyruvate is a precursor of lysine (Fig. 4A), we hypothesized that addition of pyruvate to the growth medium would counteract accumulation of (p)ppGpp due to replenishment of the lysine pool, a prediction that we indeed were able to confirm (Fig. 4B). In exponentially growing cells, 90% of CoA is present as acetyl-CoA and the synthesis and utilization of acetyl-CoA is strictly regulated (Heath & Rock, 1995). The first step of FA synthesis is catalyzed by acetyl-CoA carboxylase (ACC) that uses acetyl-CoA to make malonyl-CoA (Fig. 1A). Inhibition of FA synthesis by cerulenin (or thiolactomycin) led to an almost 100-fold increase in the malonyl-CoA pool and a concomitant 80-90% decrease in the acetyl-CoA pool within 2.5 minutes (Heath & Rock, 1995, Furukawa *et al.*, 1993). Similarly, accumulation of malonyl-CoA was also reported when cells expressing temperature sensitive enoyl-CoA reductase (FabI^ts^) were shifted to non-permissive temperature, indicating that this drastic metabolic change is common to inhibition of FA biosynthesis (Heath & Rock, 1995).

Based on our results and previous observations, we present a model explaining how FA starvation activates RelA (Fig. 6). The ACC enzyme is feedback inhibited by long-chain acyl-ACP, the terminal products of FA biosynthesis (Davis & Cronan, 2001, Parsons & Rock, 2013). Inhibition of FA chain elongation (by cerulenin or triclosan) would curtail synthesis of long-chain acyl-ACP and hence ACC will not be feedback inhibited and will keep utilizing acetyl-CoA to producing malonyl-CoA. In turn, such futile consumption of acetyl-CoA will drive continuous synthesis of acetyl-CoA from pyruvate, ultimately depleting pyruvate. Since pyruvate is an essential substrate for lysine synthesis, the low level of pyruvate would in turn curtail lysine synthesis and thereby explain the observed accumulation of uncharged tRNA^lys^ and activation of RelA (Fig. 6).

**Figure 6.**
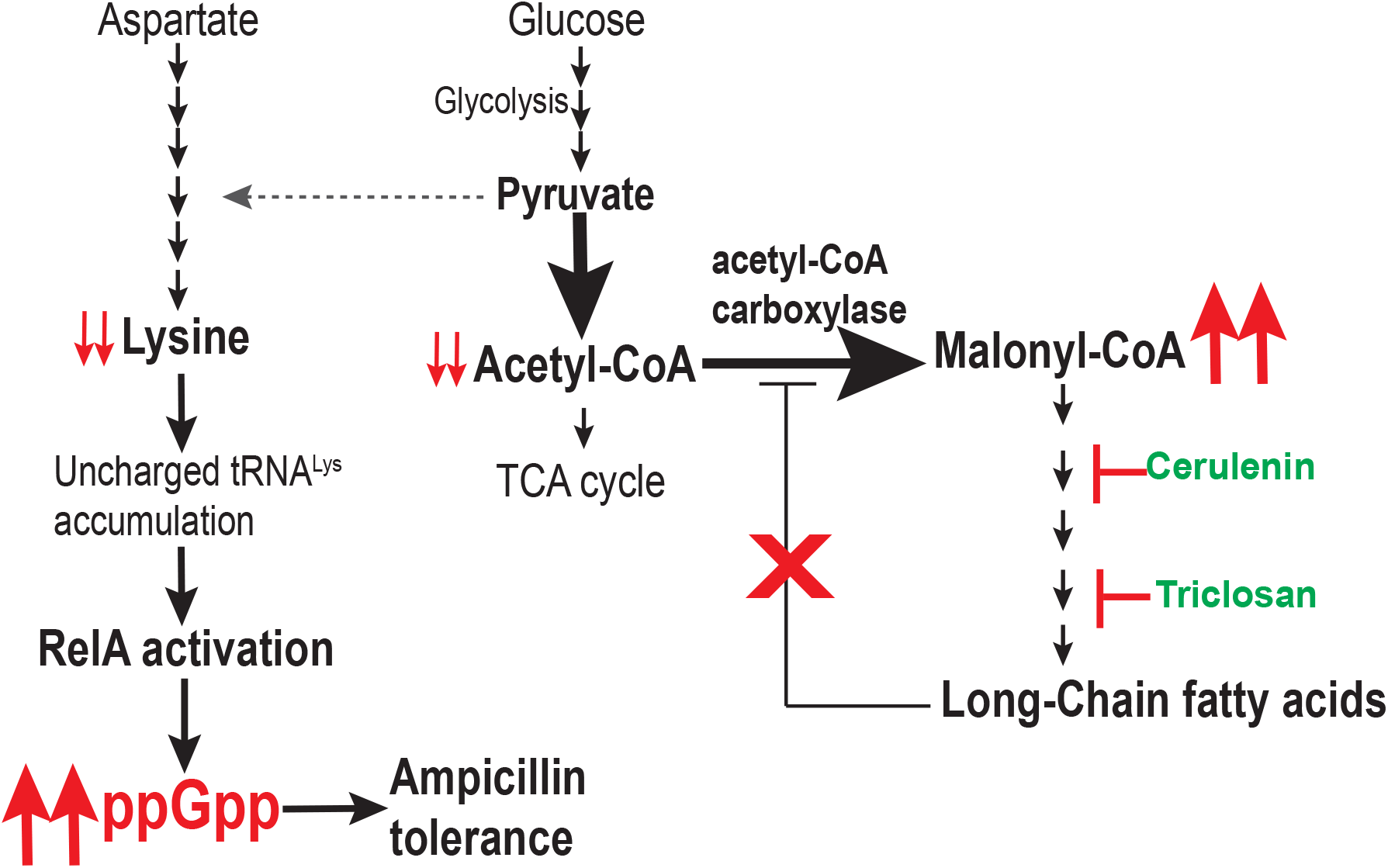
Model of RelA activation during fatty acid starvation. **(A)** During exponential growth, acetyl-CoA is synthesized and used in FA biosynthesis with strict regulation as generation of long-chain fatty acids will fine-tune the activity of acetyl-CoA carboxylase (ACC) by feed-back inhibition. This and other possible regulation mechanisms maintain very high concentration of acetyl-CoA in the cell. Acetyl-CoA is synthesized from pyruvate by the pyruvate dehydrogenase complex. Pyruvate is the resulting product of glycolysis. Pyruvate itself is required for biosynthesis of many amino acids such as lysine (shown here and in Fig. 4A), valine and isoleucine (Fig. S6C). Upon inhibition of FA biosynthesis (shown here either by cerulenin or triclosan), FA chain elongation will be blocked and long-chain FA amount drops. Low level of long-chain FA relieves the regulation of ACC causing the uninhibited reaction to proceed to consume acetyl-CoA and accumulate malonyl-CoA. We propose that this sudden change drives high rate of acetyl-CoA synthesis from pyruvate causing unavailability of pyruvate for lysine biosynthesis. Inhibition of lysine biosynthesis will deplete lysine pool and hence uncharged tRNA^lys^ levels will accumulate. RelA binds to these uncharged tRNA and vacant A-site on ribosome to synthesize ppGpp. High ppGpp levels shut down cell wall synthesis resulting in ampicillin tolerance. Red color shows the consequences of FA starvation; Thick arrow shows enhanced activity of enzyme/ accumulation of products; dotted arrow shows reduced activity of enzyme/depletion of products.

Thus, pyruvate appears to be the connecting link between FA starvation and lysine starvation. Pyruvate plays an important role in general metabolism. For instance, many amino acids directly or indirectly use pyruvate for their synthesis; and acetyl-CoA synthesized from pyruvate can either be used for FA synthesis or enter the TCA cycle. Using *E. coli* cell extracts, it was shown that pyruvate supplementation partially relieves valine-induced inhibition of isoleucine biosynthesis, since pyruvate is one of the substrates in the reaction inhibited by valine (Fig. S6C) (Leavitt & Umbarger, 1962). However, an excess of pyruvate did not cause a significant reduction in ppGpp accumulation upon valine treatment (Fig. S6B). Considering the importance of pyruvate, it is understandable that many genes encoding enzymes involved in pyruvate metabolism are upregulated during the stringent response with an overall effort of the cell to maintain pyruvate at a sustainable level (Traxler *et al.*, 2008).

Lysine starvation is the result of FA starvation, which raises new important questions regarding our understanding of RelA function and the stringent response under conditions that are not directly linked to amino acid starvation. Apart from FA starvation, glucose starvation and phosphate starvation have also been shown to induce a similar higher and rapid ppGpp accumulation in *wt* compared to cells lacking RelA, suggesting a much broader role of RelA as perhaps the major (p)ppGpp synthetase (Gentry & Cashel, 1996, Spira *et al.*, 1995, Murray *et al.*, 2003). SpoT has been described to sense FA, glucose, phosphate, and iron starvation and can synthesize alarmone in cells lacking RelA, but how its activity will be regulated when RelA is present to quickly synthesize high amount of (p)ppGpp, will be an important question to address. The paralogous *relA* and *spoT* genes are derived from a common ancestor, the bifunctional *rel*/*rsh* gene by duplication in proteobacteria and sequence analysis of SpoT in the Moraxellaceae family of gammaproteobacteria suggests that its synthetase activity has been gradually weakened during evolution (Mittenhuber, 2001, Atkinson *et al.*, 2011).

FA biosynthesis in bacteria has been an attractive target for new antibiotics as bacterial FA biosynthesis enzymes are structurally different from their eukaryotic analogues. However, increasing evidence suggests that high concentration of FA inhibitors induce survival and cross-resistance to other antibiotics (Rodionov & Ishiguro, 1996, Westfall *et al.*, 2019, Kampf, 2018, Fahimipour *et al.*, 2018). Many of these inhibitors are being used in products ranging from animal feed to personal hygiene products such as hand wash, disinfectants, and get released to the environment. Here bacteria will be exposed to sub inhibitory concentrations of the drugs, which can have a wide range of consequences. In this study, we report here that exposure to low concentrations of FA inhibitors even for a short period of time (5 minutes), is sufficient to induce RelA mediated stringent response (Fig. 1) and provide a 100-fold higher tolerance to cell wall targeting antibiotics (Fig. 5A to 5C). Recently, RelA dependent antibiotic tolerance was also observed during nitrogen starvation (Brown, 2019). Interestingly, the observation that the antibiotic tolerance induced by FA inhibitors can be completely reversed by supplementing lysine shows therapeutic potential of our study (Fig. 5A to 5C). Interestingly, at high doses of cerulenin (~4X MIC) the cells become highly tolerant to ampicillin lysis independently of *relA*. This effect could be due either to SpoT mediated ppGpp synthesis and/or by complete shutdown of phospholipid biosynthesis, as suggested previously (Rodionov & Ishiguro, 1996).

Taken together, our data robustly show that in *wt* cells, RelA is the dominant ppGpp synthetase under FA starvation by sensing lysine starvation arising because of pyruvate depletion. We propose that other starvation conditions may lead to amino acid starvation in turn raising the possibility that there is a functional segregation such that RelA has become the dominant synthetase and SpoT the only (p)ppGpp hydrolase of *E. coli*. This line of reasoning is supported by the observation that the initial burst of ppGpp synthesis during glucose starvation depends on RelA and is suppressed by the addition of amino acids to the growth medium (Gentry & Cashel, 1996). Our work sheds new light on how the stringent response is triggered and will provide a novel perspective of how to analyze the response under various stress conditions.

## Experimental Procedures

All growth media supplements such as MOPS, glucose, amino acids and other chemicals such as ampicillin, cerulenin, and triclosan were purchased from Sigma. γ-[^32^P]-ATP was obtained from PerkinElmer, USA and [^32^P] phosphoric acid was from Hartmann Analytic, Germany.

### Strains

MG1655*ΔldcCΔcadA* was created by P1 transduction of *ΔldcC::kan* and *ΔcadA::kan* of the KEIO collection strains (Baba *et al.*, 2006) into MG1655 to generate MG1655*ldcC cadA*. After each transduction, the *kan* resistance cassette was removed using the FLP recombinase provided from plasmid pCP20 (Datsenko & Wanner, 2000). Correct resistance cassette insertion and removal were confirmed by PCR. The resulting strain lacks *ldcC* and *cadA* encoding lysine decarboxylases.

### Media and growth conditions

The standard *Escherichia coli* K-12 strain MG1655 (*wt*) strain and its *ΔrelA* derivative were from the lab collection. Cells were routinely streaked from frozen stock onto LB plates and incubated at 37°C overnight for revival. For all experiments, single colonies from the freshly streaked plate were inoculated and grown overnight at 37°C in MOPS minimal media supplemented with 0.2 % glucose, all nucleobases, and 1.32 mM K_2_HPO_4_ as described earlier (Winther *et al.*, 2018, Neidhardt *et al.*, 1974). Culture of *ΔrelA* cells were routinely checked for their growth sensitivity on SMG plates to rule out for the presence of any suppressor mutations.

### Minimum inhibitory concentration (MIC) calculation

MIC calculation was according to (Wiegand *et al.*, 2008). Briefly, overnight inoculates were grown at 37°C in MOPS minimal medium supplemented with 0.2 % glucose and diluted to an OD_600_ of ~0.01 and grown to ~0.2 and again diluted to ~0.01. Drugs to be tested (cerulenin and triclosan) were serially diluted into the same MOPS glucose growth medium. A volume of 150 µl of a culture at OD_600_ of ~0.01 was mixed with 150 µl serially diluted drug to obtain a final concentration half of the initial concentration of the drug. They were placed in microtitre plates and OD_600_ was monitored every 15 minutes with shaking in a BioTek microplate reader for 12-14 hrs. MIC was determined as the lowest concentration at which no growth observed after 12 hrs of incubation.

### Measurement of cellular levels of (p)ppGpp

Estimation of [(p)ppGpp] was done as described previously with a few modifications (*Winther et al., 2018*). Briefly, overnight inoculates were grown at 37°C in high phosphate (1.32 mM K_2_HPO_4_) MOPS minimal medium supplemented with 0.2 % glucose (or 0.5 % glycerol as mentioned) (Neidhardt *et al.*, 1974). Cultures were then diluted to 0.01 OD_600_ in low phosphate (0.2 mM K_2_HPO_4_) MOPS glucose minimal medium and grown till 0.3 OD_600_. At 0.3 OD_600_, they were again diluted 10-fold in the same growth medium supplemented with ^32^P (final specific activity 100 µCi/ml) and grown for 2-3 generations. Fatty acid starvation was induced either by the addition of cerulenin or triclosan, as mentioned for the individual experiments. Amino acid starvation was induced by addition of valine (500 µg/ml). Fifty-microliter samples were withdrawn and mixed with 10 µl of ice-cold 2 M formic acid before and 5, 10, 15, 20 and 30 minutes after starvation. After two freeze-thaw cycles, samples were centrifuged at 4°C to pellet the cell debris, and 5 µl aliquots of the supernatants were applied to polyethyleneimine (PEI) cellulose thin layer chromatography plates (Merck, Sigma), resolved with 1.5 M KH_2_PO_4_ (pH 3.4) air dried and exposed by phosphorimaging (Amersham Typhoon phosphorimager). Quantification was done using ImageJ and relative aboundance of ppGpp and pppGpp was estimated as a percentage value relative to the total (pppGpp+ppGpp+GTP) amount as described (Fernandez-Coll & Cashel, 2018, Mechold *et al.*, 2013).

### Pyruvate quantification assay

Cellular pyruvate level was quantified using Pyruvate Assay Kit (Sigma) according to manufacturer’s protocol. Briefly, cells were grown in balanced growth in MOPS glucose media as above. Cells were collected before and 10 minutes after treatment with triclosan (1 µg/ml). Cells were immediately put on ice for 15 minutes and then pelleted down in precooled centrifuge and resuspended in pyruvate assay buffer. Cells were homogenized using glass beads (Sigma) in homogenozer MagNA Lyser (Roche). Centrifuged for 13000 X g for 10 minutes to remove insoluble material. Samples (50 µl) were used for fluorometric pyruvate assay and fluorescence intensity (λ_ex_=535/λ_em_ =587 nm) was measured as per protocol.

### Ampicillin tolerance assay

In order to determine FA starvation dependent survival to ampicillin exposure, single colony of *wt* and *ΔrelA* cells were inoculated for overnight growth at 37 °C in MOPS minimal medium supplemented with 0.2 % glucose as described above (Neidhardt *et al.*, 1974). The cells were then diluted to an OD_600_-0.01 in the same media and grown till 0.3 OD_600_. To keep cultures in balanced growth, cells were again back-diluted to 0.03 OD_600_ and grown till ~0.1 OD_600_. At this point, 0.4 mM lysine was added in one aliquote of *wt* cells and grown for 5-10 minutes. Aliquots of *wt*, *relA* and *wt* (+lysine) were treated with cerulenin (50 or 200 µg/ml) or triclosan (1 µg/ml) (or valine (500 µg/ml) for separate experiments) for 5 minutes and then exposed to ampicillin (100 µg/ml) for 3 to 4 hrs while shaking. One aliquot of each culture was directly exposed to ampicillin without being pre-exposed to cerulenin or triclosan (or valine). Untreated samples were collected before ampicillin exposure. After ampicillin exposure, cells were centrifuged, supernatant containing ampicillin was discarded, washed with sterile phosphate-buffered saline (PBS) and cell pellets were resuspended in fresh medium to concentrate five times. Samples were serially diluted and 5 µl of each dilution was spotted on LB plates, dried and incubated overnight at 37°C incubator to quantify survivors.

### tRNA charging levels

tRNA charging levels were analysed using Northern blotting method as described earlier (Winther *et al.*, 2018). Briefly, bacterial cultures were grown overnight in MOPS minimal medium supplemented with 0.2% glucose at 37°C. Next morning they are diluted to 0.01 OD_600_ and grown till ~0.2 OD_600_. 4mL sample was collected as before starvation and mixed with 4 mL of cold 10% TCA and incubated on ice. Triclosan (1µg/ml) was added into the remaining culture to induce fatty acid starvation. Samples were collected at different time points. For lysine supplementation, similar method was used but 0.4 mM lysine was added 10 minutes before triclosan addition into the culture. tRNAs were isolated, separated by denaturing PAGE, transferred to a Hybond N+ membrane and hybridization were performed using the same protocol as described earlier (Winther *et al.*, 2018). Lysine tRNA (Lys) and serine tRNA (Ser3) probes were synthesized using published sequences (Dong *et al.*, 1996).

## Acknowledgements

We thank Michael A. Sørensen and Sine Lo Svenningsen for discussion and suggestions. We would also like to thank Sidsel K. Henriksen for efficient management of lab and making quick availability of reagents and chemicals. This project was funded by grants from the Novo Nordisk Foundation (to K.G) and the Danish National Research Foundation (DNRF 120).

---

**Figure S1.**
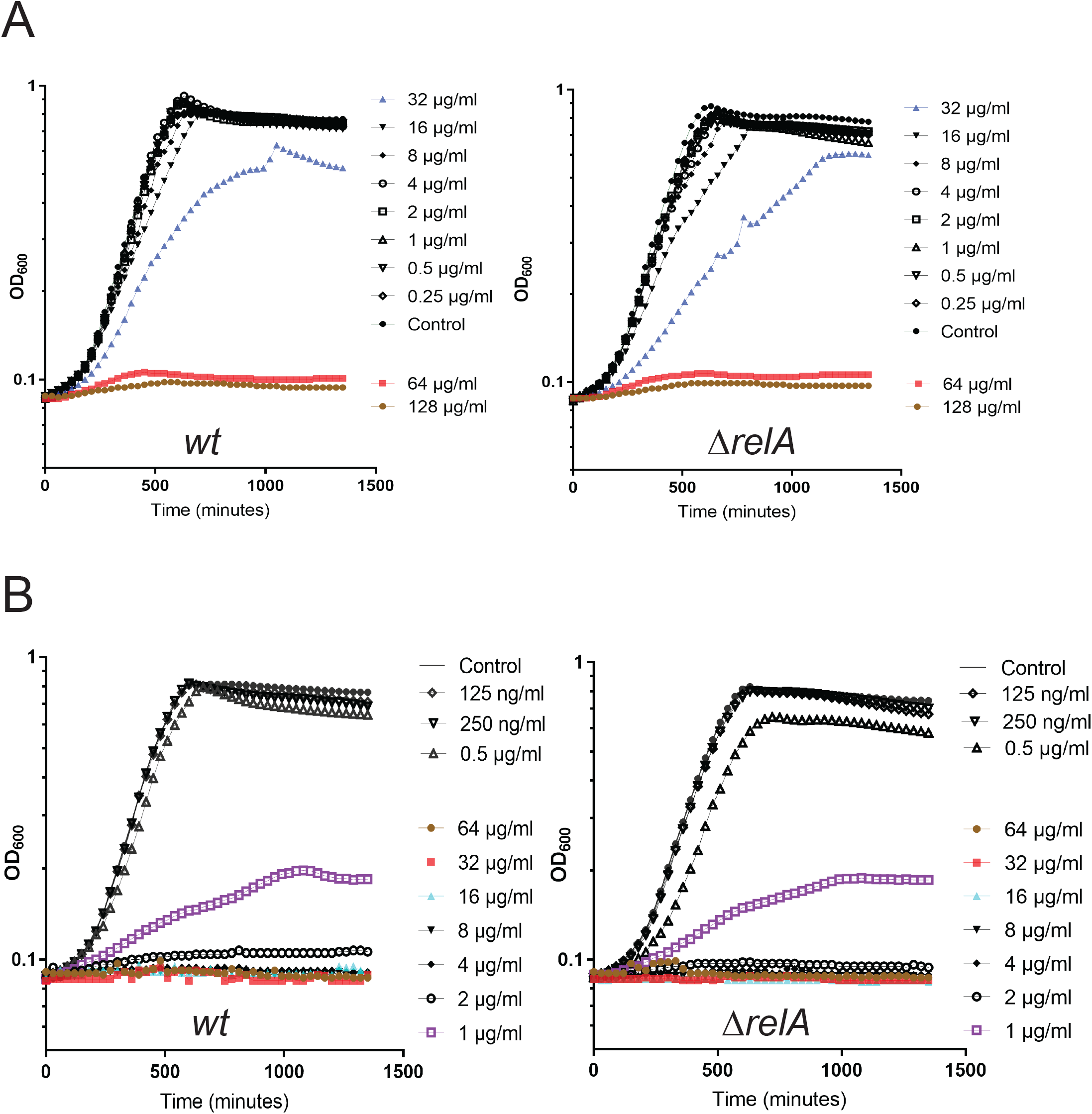
Minimum inhibitory concentration of cerulenin and triclosan for *wt* and Δ*relA* cells. **(A)** MIC of cerulenin was calculated for *wt* and Δ*relA* as described in materials and methods. At 64 µg/ml final concentration of cerulenin no growth was observed in *wt* and in Δ*relA* cells therefore 64µg/ml was considered as MIC value for both the strains. **(B)** MIC of triclosan was calculated for *wt* and Δ*relA* as described in materials and methods. At 1 µg/ml final concentration of triclosan little growth was observed either in *wt* or in Δ*relA* cells therefore 1µg/ml was considered as MIC value of triclosan for both the strains. MIC for both *wt* and Δ*relA* cells was found to be close to ~1 µg/ml for triclosan were no growth observed after 24 hrs of incubation in MOPS glucose medium (Fig. S1B).

**Figure S2.**
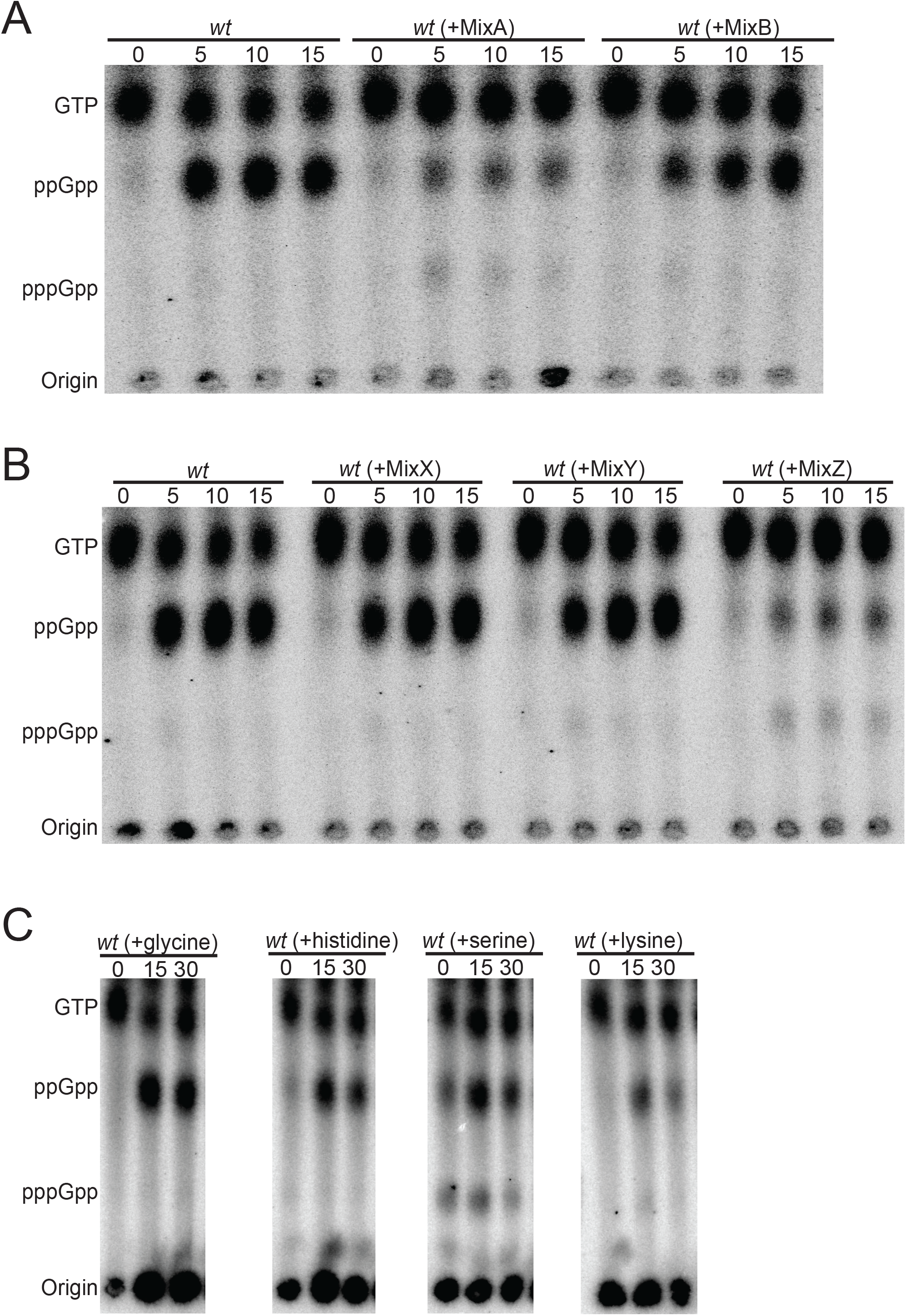
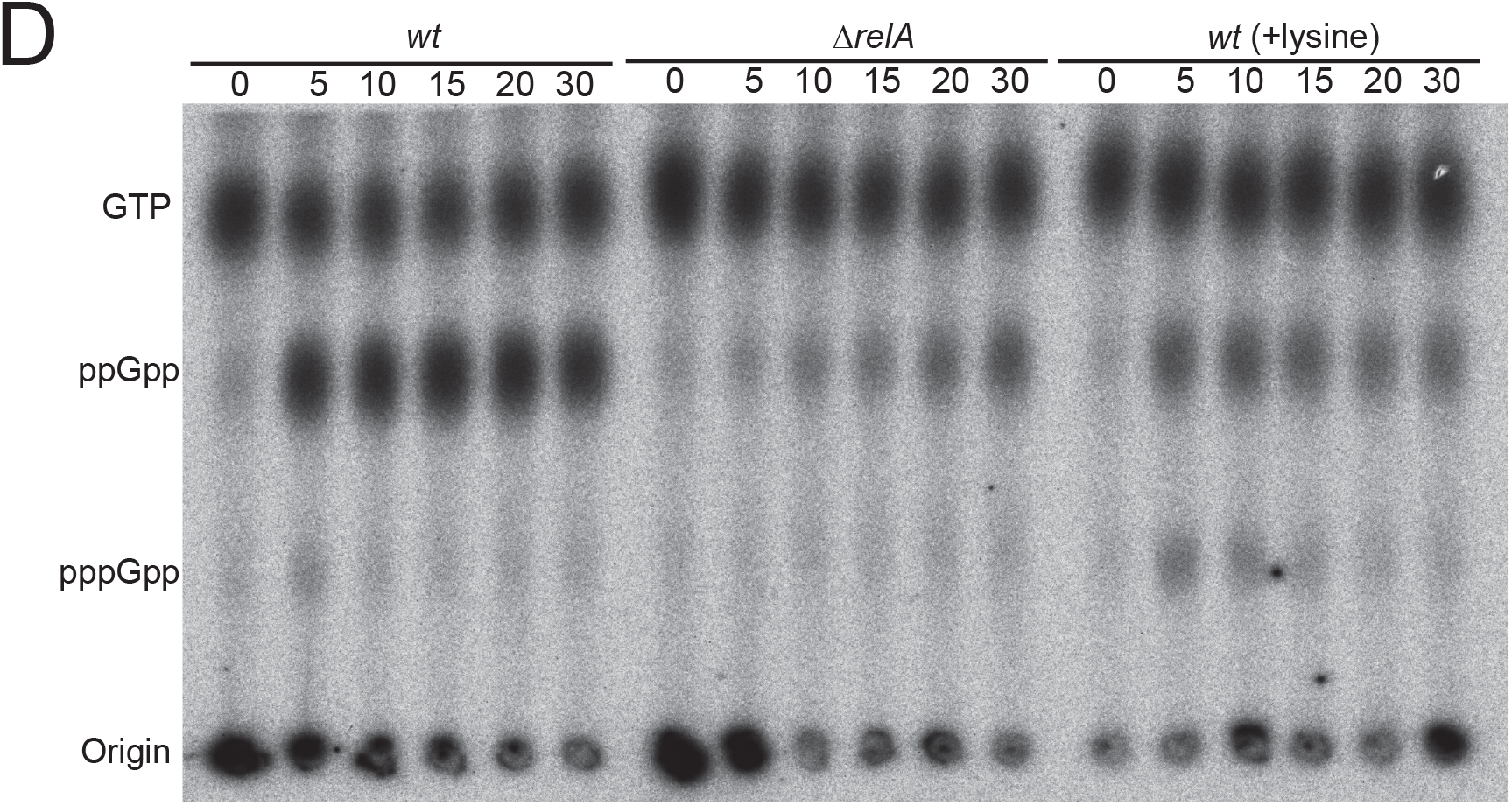
Effect of amino acid mixture supplementation on stringent response during fatty acid (FA) starvation. **(A)** Representative autoradiogram of a TLC showing the time dependent (p)ppGpp accumulation upon cerulenin (200 µg/ml) treatment in *wt* cells either without or pretreated with mixture of 10 amino acids each in MixA and in MixB. MixA (alanine, arginine, asparagine, aspartate, glutamate, glutamine, glycine, histidine, serine and lysine) and MixB (isoleucine, proline, tyrosine, threonine, tryptophan, valine, leucine, cysteine, methionine and phenylalanine) **(B)** As in A but cultures were pretreated with mixture of amino acids from MixX, MixY or MixZ. MixX (alanine, arginine, asparagine), MixY (aspartate, glutamate, glutamine) and MixZ (glycine, histidine, serine and lysine). **(C)** As in A, but cultures were pretreated with single amino acids glycine, histidine, serine or lysine. All amino acids were used at final concentration as recommended (Neidhardt *et al.*, 1974). Positions of GTP, ppGpp and pppGpp are indicated. Time points above TLC images are in minutes after cerulenin treatment. **(D)** Representative TLC showing the time dependent (p)ppGpp accumulation upon cerulenin (200 µg/ml) treatment in *wt*, Δ*relA* or *wt* (supplementated with 0.4 mM lysine).

**Figure S3.**
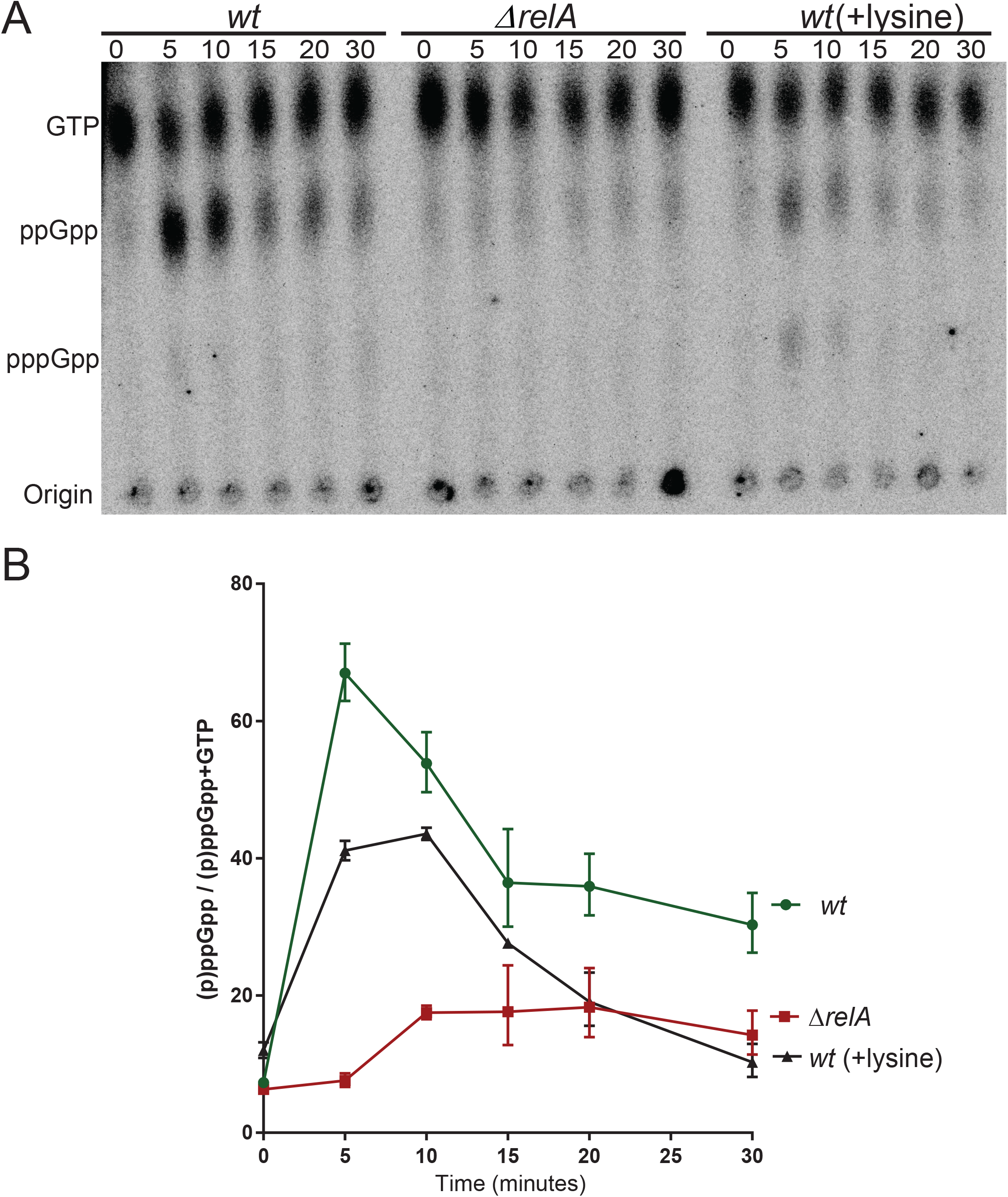
Lysine supplementation also reduces stringent response during triclosan treatment. **(A)** Representative TLC image showing the time dependent (p)ppGpp accumulation upon triclosan (1 µg/ml) treatment in *wt*, *ΔrelA* and *wt* (supplemented with 0.4 mM lysine). **(B)** Quantification of (p)ppGpp accumulation in *wt*, *ΔrelA* and *wt* (supplemented with 0.4 mM lysine) upon triclosan (1 µg/ml) treatment. Data collected from three different experiments were used to calculate mean with standard deviation.

**Figure S4.**
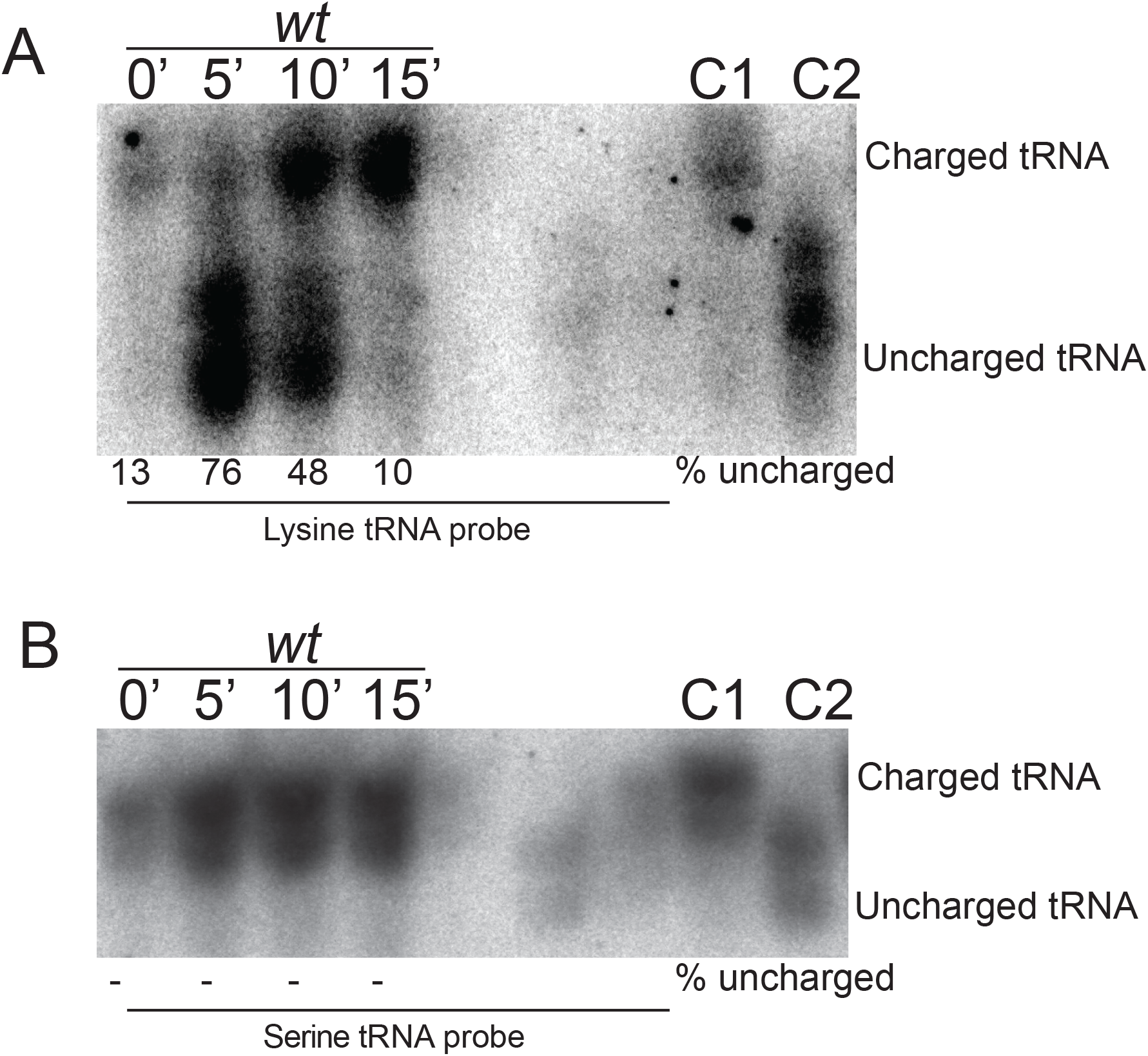
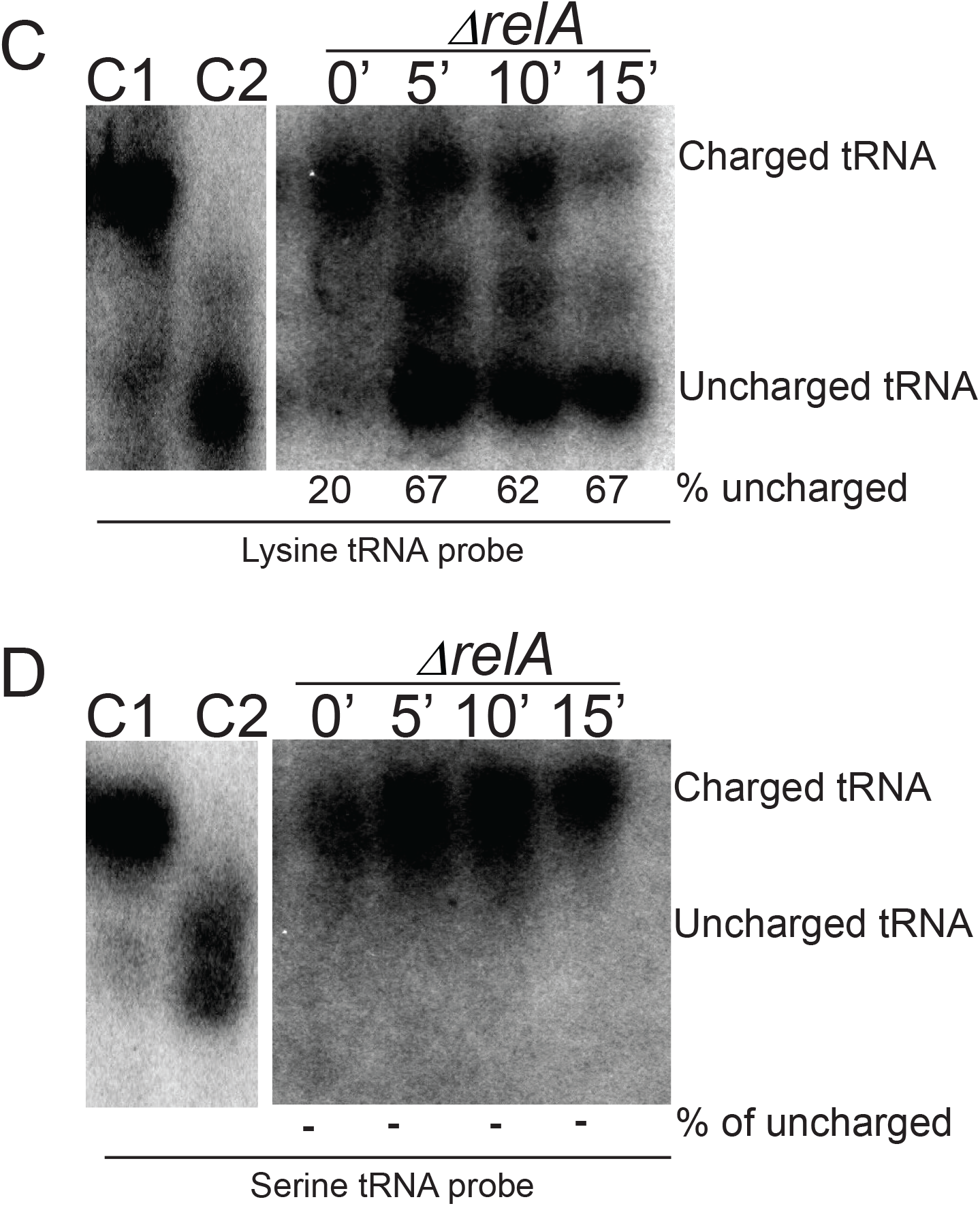
tRNA acylation levels of *wt* and Δ*relA* cells before and after triclosan (1µg/ml) treatment. Total tRNA isolated from *wt* **(A)** or ∆*relA* **(C)** cells before and 5, 10, 15 minutes after triclosan (1µg/ml) treatment and processed for Northern blotting analysis using an probe specific to tRNAlys. Percentage (%) of uncharged RNA was quantified according to signal intensity of uncharged/(charged+uncharged). A fraction of tRNA was deacylated in-vitro by base treatment (100 mM Tris-HCl [pH-9]) for control. Positions of charged (C1) and uncharged (C2) controls are shown. **(B)** & **(D)** Blots in (A) & (C) are hybridized with tRNAser specific probe repectively.

**Figure S5.**
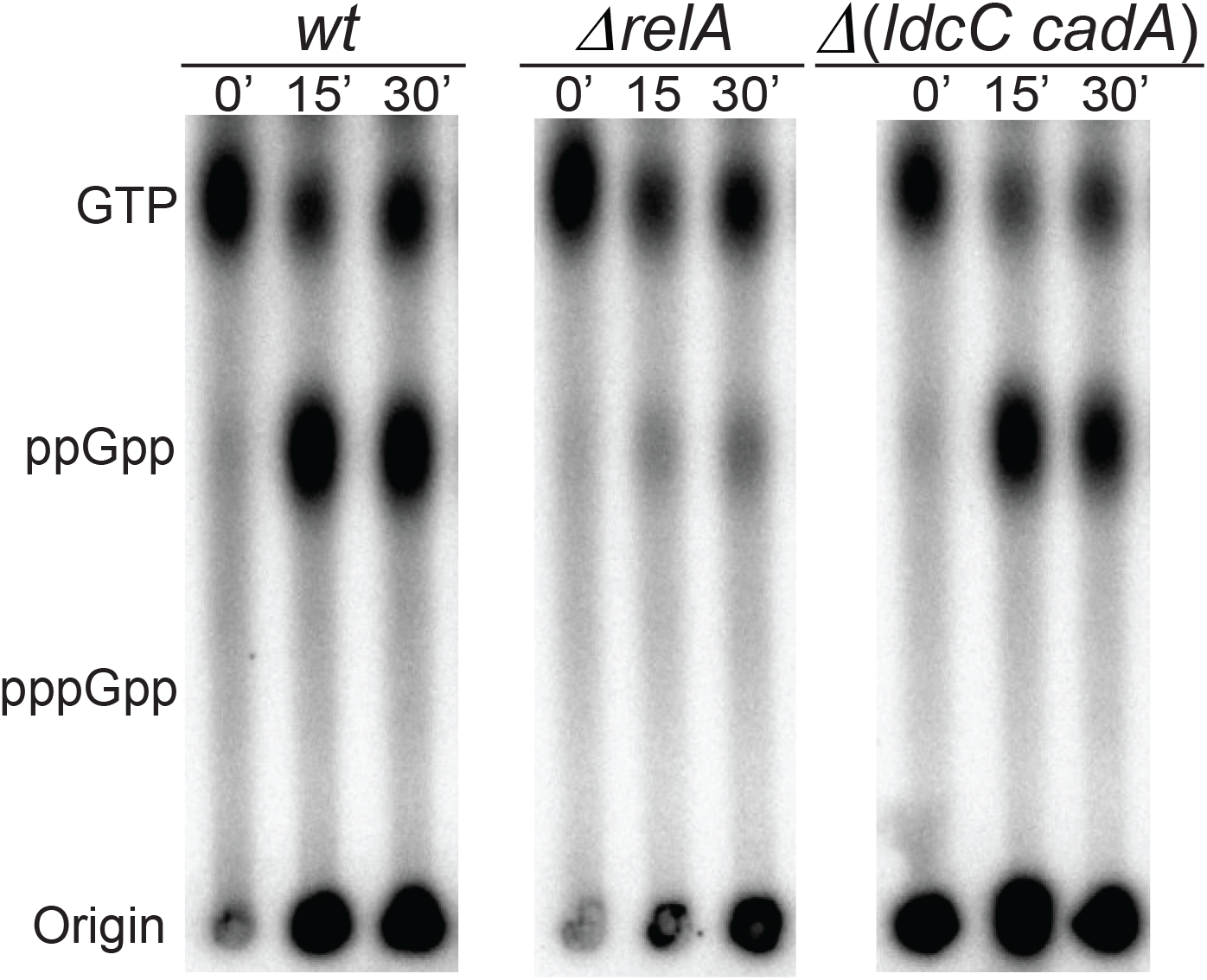
Lysine decarboxylase mutant does not affect the stringent response after fatty acid starvation. Representative TLC image showing the time dependent (p)ppGpp accumulation upon cerulenin (200 µg/ml) treatment in *wt*, Δ*relA* and in Δ*ldcC* Δ*cadA* mutant. The positions of GTP, ppGpp and pppGpp are indicated. Time-points above TLCs are in minutes after cerulenin treatment. Levels of (p)ppGpp were measured as described in the Legend to Fig. 1.

**Figure S6.**
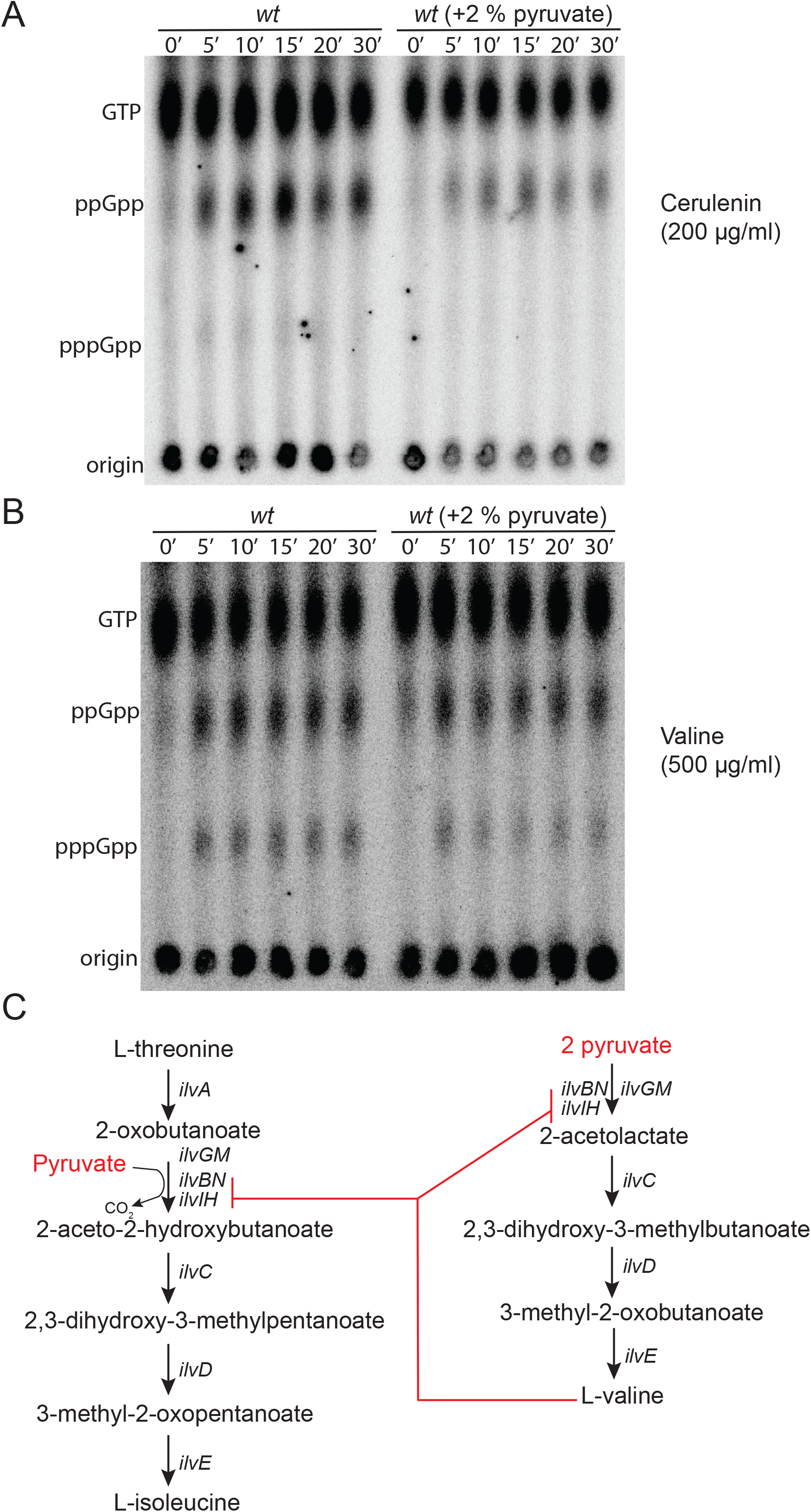
Effect of pyruvate supplementation on stringent response. **(A)** (p)ppGpp accumulation upon cerulenin (200 µg/ml) treatment in *wt* with and without pretreatment with 2% pyruvate. Levels of (p)ppGpp were measured as described in the Legend to Fig. 1. **(B)** Representative TLC image showing the time dependent (p)ppGpp accumulation upon valine (500 µg/ml) treatment in *wt* (untreated) and when pretreated with 2% pyruvate. Positions of GTP, ppGpp and pppGpp are indicated. Time points above TLC images are in minutes after cerulenin or valine treatment. Levels of (p)ppGpp were measured as described in the Legend to Fig. 1. **(C)** Schematic representation of valine and isoleucine biosynthesis pathways in *E. coli* K-12 (Keseler *et al.*, 2017). Excess valine inhibits enzymes that use pyruvate as one of the substrates (shown in red). Pyruvate supplementation has previously been shown to relax this inhibition in a competitive manner (Leavitt & Umbarger, 1962).

**Figure S7.**
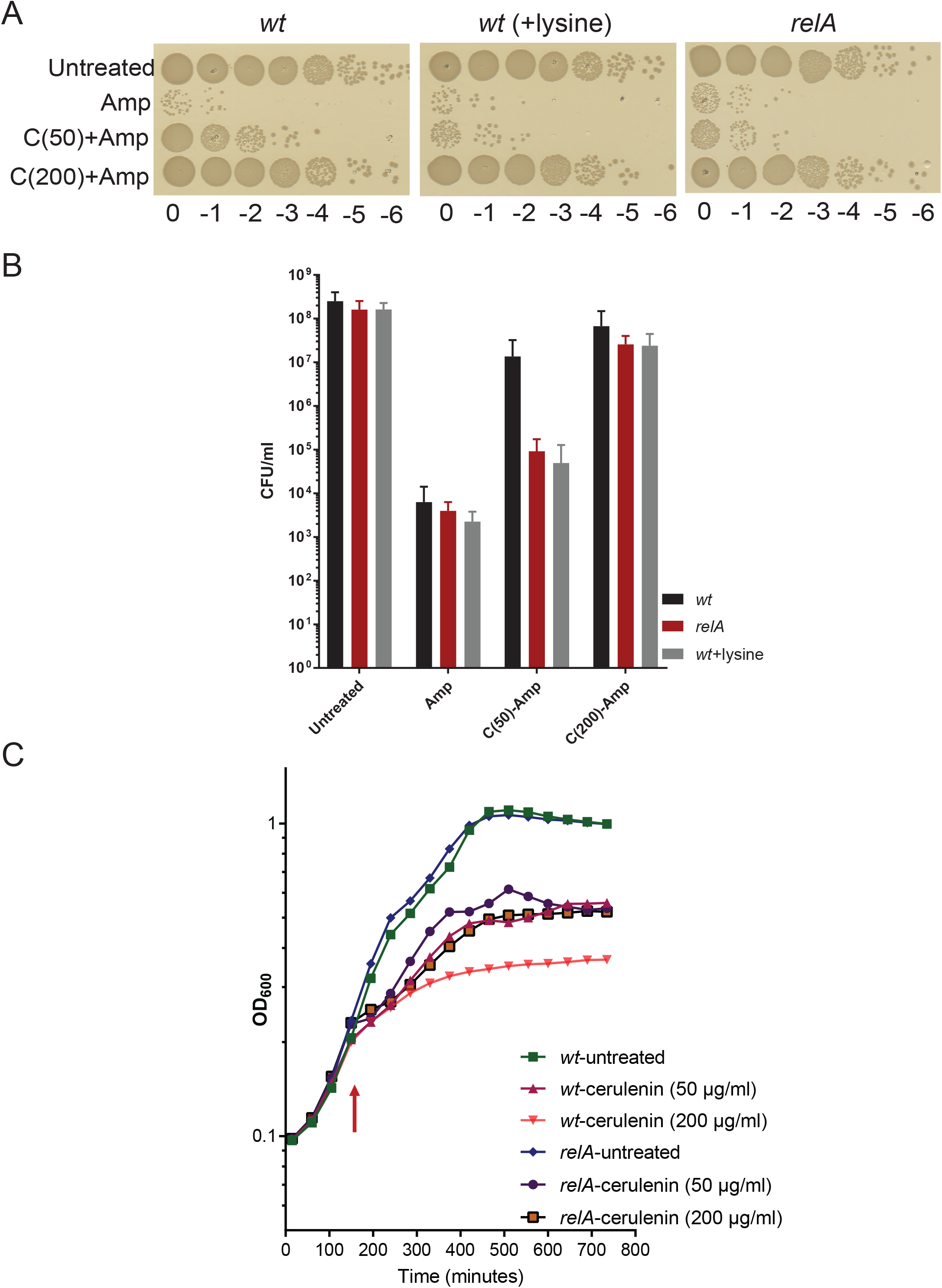
Fatty acid starvation increases tolerance to ampicillin. **(A)** *wt* and Δ*relA* cells growing balanced in MOPS + 0.2 % glucose media were either left untreated, directly exposed to ampicillin (100 µg/ml) for 3 hrs or first treated with cerulenin (either 50 or 200 µg/ml) for 5 minutes and then exposed to ampicillin. In another culture of the *wt* strain, 0.4 mM lysine was added and then processed in the same way to analyse the effect of lysine on ampicillin tolerance. **(B)** Colony forming units (CFU) from four different experiments, performed as in A. Means and standard deviations are indicated. **(C)** Cells of *wt* and Δ*relA* strains growing balanced in MOPS + 0.2 % glucose medium measured in a plate reader up to OD_600_ ~0.2 and then cerulenin was added (50 or 200 µg/ml). Arrow indicates addition of cerulenin. Growth rate was monitored at 15’ intervals for 12-14 hrs. Data points are averages of four experiments.

